# Protection against doxorubicin-induced cardiotoxicity by ergothioneine

**DOI:** 10.1101/2022.12.21.521347

**Authors:** Irwin K. Cheah, Richard M.Y. Tang, Xiaoyuan Wang, Karishma Sachaphibulkij, Suet Yen Chong, Lina H.K. Lim, Jiong-Wei Wang, Barry Halliwell

**Author notes:** Co-corresponding authors: Prof. Barry Halliwell, National University of Singapore, Life Science Institute, Neurobiology Programme, Centre for Life Sciences, 28 Medical Drive, #05-01, Singapore 117456, Email address, Dr. Jiong-Wei Wang, National University of Singapore Cardiovascular Research Institute, 14 Medical Drive, 08-01, Singapore 117599. Authors contributed equally to this work (Irwin K. Cheah, Richard M.Y. Tang).

## Abstract

Anthracyclines such as doxorubicin remain the first line of treatment for haematological malignancies, and breast cancers. However, the potential risk of cardiac injury by anthracyclines, which may lead to severe myopathy or heart failure, severely limits their application, and remains a challenge to ensuring curative chemotherapy. While the complex interplay between pathological pathways of anthracycline cardiotoxicity is yet to be fully understood, oxidative damage, iron overload-mediated formation of reactive oxygen species (ROS), mitochondrial dysfunction, and inflammation are all believed to be involved.

The unique dietary thione, ergothioneine, while not produced in animals and humans, can be avidly absorbed and accumulated in tissues including the heart. Amongst other cytoprotective properties ergothioneine has been shown to scavenge various ROS, decrease proinflammatory mediators, chelate metal cations such as Fe^2+^, preventing them from partaking in redox activities, and may protect against mitochondrial damage and dysfunction. Moreover, low plasma ergothioneine levels are also strongly correlated to risk of cardiovascular events in humans, suggestive of a cardioprotective role. Taken together this highlights the potential for ergothioneine to counteract anthracycline cardiotoxicity.

Here we investigate the potential of ergothioneine supplementation to protect against cardiac dysfunction in mice models of doxorubicin-induced cardiotoxicity and found that it had significant protective effects. Moreover, ergothioneine administration in a mouse breast cancer model did not exacerbate the growth of the tumour and did not interfere with the chemotherapeutic efficacy of doxorubicin. These results suggest that ergothioneine could be a viable co-therapy to alleviate the cardiotoxic effects of anthracyclines in the treatment of cancers.

## Introduction

Anthracyclines, such as doxorubicin, are widely used chemotherapy drugs in both adults and children for haematologic malignancies, soft-tissue sarcomas, and solid tumours of the breast, lung, stomach, bladder, and ovaries. Despite their use as a standard for effective curative chemotherapy for decades, their clinical application can be severely limited due to dose-specific cardiotoxic side effects, which range from asymptomatic electrocardiographic changes to severe cardiomyopathy and heart failure [1]. These cardiotoxic effects can persist long after cessation of chemotherapy, and in some cases myocardial damage may not be apparent until many years after completion of chemotherapy and can be exacerbated by other cardiometabolic co-morbidities, such as hypertension, coronary artery disease and diabetes [2]. Doxorubicin, one of the most effective anthracycline drugs, is frequently prescribed but it also poses the greatest risk of cardiac injury. It is estimated that over one quarter of patients receiving a cumulative dose of doxorubicin exceeding 550 mg/m^2^ will develop heart failure [2]. Despite decades of research and increased understanding of anthracycline cardiotoxicity, maintaining chemotherapeutic efficacy while preventing cardiotoxicity remains a challenge, especially in children [1].

While the exact pathological mechanisms of anthracycline cardiotoxicity are not fully understood, it is known to involve multiple pathways including excessive oxidative stress [3], activation of pro-inflammatory pathways [4], mitochondrial dysfunction [5], mitochondrial iron overload-induced lipid peroxidation [6], and activation of apoptotic pathways [7]. Studies suggest a strong affinity of anthracyclines to cardiolipin, a phospholipid found abundantly on the inner mitochondrial membrane, explaining the tendency for doxorubicin and its complexes with iron to accumulate in the mitochondria [8]. Since cardiomyocytes are rich in mitochondria, to meet their high energy demands, this may explain the predisposition of the heart to damage by anthracyclines. Redox cycling by doxorubicin involving mitochondrial complex I [3] together with mitochondrial iron overload [9] leads to the formation of reactive oxygen species (ROS) such as superoxide and hydroxyl radials, and hydrogen peroxide. Together with anthracycline-induced downregulation of glutathione peroxidase 4, this contributes to excessive lipid peroxidation and ultimately cardiomyocyte ferroptosis; a process which has been shown to be mitigated by iron chelators [6].

The maintenance of redox homeostasis is critical to the preservation of normal cardiomyocyte function [10] and disruption of this by anthracyclines through pro-oxidative mechanisms mentioned earlier can disrupt redox sensitive signalling pathways and damage important cellular components, ultimately causing cardiomyocyte death [11]. As oxidative stress has been hypothesised to be one of the primary drivers of anthracycline cardiotoxicity, some studies have evaluated the application of antioxidant compounds, such as vitamins C [12], and E [13], and N-acetylcysteine [14], to prevent cardiac injury. While outcomes are somewhat mixed, a few studies have yielded positive outcomes *in vitro* and in acute toxicity *in vivo* models, however none have proven successful in a clinical setting [15, 16]. Moreover, there is concern as to whether these compounds may interfere with the chemotherapy [17, 18], exacerbate tumour growth, or promote metastasis as has been suggested by a few animal studies (e.g. [19-21]). At present, the iron chelator, dexrazoxane, is the only drug approved by the US FDA to counteract anthracycline cardiotoxicity. It appears to work by binding iron from the anthracycline-iron complexes and inhibiting formation of ROS [22]. However, it does not eliminate cardiotoxic risk and there are concerns of secondary haematological malignancies and reduced response to chemotherapy, which is possibly why dexrazoxane is only prescribed to a small percentage of patients undergoing doxorubicin chemotherapy [23].

The effectiveness and widespread application of anthracyclines in cancer therapy means their removal is not a viable solution, hence the absence of an effective prophylactic cardioprotectant against anthracyclines highlights an important therapeutic gap. Prior studies have shown that a low molecular weight dietary thione, ergothioneine (ET), may act as a physiological cytoprotectant (for detailed reviews on this compound please see [24-27]). ET is not known to be produced in animals and humans but is abundant in diets, especially mushrooms, and is avidly taken up and accumulated in tissues including the heart [25, 28], by the cation transporter, OCTN1, also known as the ET transporter due to its specificity for ET [29, 30]. Many studies have highlighted the cytoprotective properties of ET including the ability to scavenge free radicals [31], decrease pro-inflammatory mediators, chelate divalent metal cations (such as Fe^2+^), preventing their redox activities [31-33], and prevent mitochondrial damage and dysfunction [34, 35] suggesting that it could be a cardioprotective agent against anthracyclines. Indeed higher levels of ET in human plasma were identified as an independent marker of lower risk of cardiometabolic disease and associated mortality in a longitudinal study involving 3236 participants [36]. Other studies have also demonstrated that ET is able to protect the vascular system against diabetes-induced cardiovascular injury in rats [37], protect the endothelium against a range of toxins to prevent endothelial dysfunction [38-40], and protect tissues against ischaemia-reperfusion injury [41-43]. Moreover, ET reduced levels of proinflammatory cytokines [43, 44] and expression of vascular adhesion molecules (VCAM-1, ICAM-1 and E-selectin) [45], and was shown to reduce the pro-oxidant ferrylmyoglobin to metmyoglobin [46], highlighting possible ways that ET could protect the heart from inflammatory and oxidative injury induced by anthracyclines. Untargeted metabolomic studies revealed that levels of ET were elevated in the heart of mice subjected to either transverse aortic constriction (pressure overload) or myocardial infarction [47], supporting our prior hypothesis [48] that tissues may upregulate expression of the ET transporter to increase ET uptake in injured tissue as a cytoprotective mechanism [49].

This evidence suggest that ET could be an excellent candidate to counteract multiple pathological mechanisms of anthracycline cardiotoxicity, however to date this has not been investigated. The present study demonstrates that ET is able to improve cardiac function in a mouse model of doxorubicin-induced cardiotoxicity. Moreover, in a breast cancer model, supplementation of ET in the presence or absence of doxorubicin administration demonstrated that ET neither exacerbated tumour growth nor interfered with the chemotherapeutic efficacy of doxorubicin. This study presents a strong case for ET to be used in conjunction with anthracycline chemotherapy to counter cardiac injury and lays the foundations for future clinical evaluation of ET against anthracycline cardiotoxicity.

## Methodology

### Chemicals and reagents

L-ergothioneine (>98%), L-ergothioneine-d9 (ET-d_9_), hercynine, and hercynine-d_9_, were provided by Tetrahedron (Paris, France). Doxorubicin, hydrochloride salt (>99%) was purchased from LC Laboratories (MA, USA). All other chemicals were purchased from Sigma-Aldrich (MO, USA), unless otherwise stated.

### Animal studies

Mice (C57BL-6J, BALB-C) were randomly assigned to cages in groups of 4-5 animals and provided with water and standard mice chow (Teklad, USA) *ad libitum*. The animals were kept in a barrier animal facility with a 12h/12h light-dark cycle. Animals were cared for in accordance with the protocols and guidelines stipulated in the Institutional Animal Care and Use Committee (IACUC), National University of Singapore. All the procedures were performed with prior approval by the IACUC, NUS; protocol no. R17-1242.

### Doxorubicin cardiotoxicity model

To induce anthracycline cardiotoxicity, C57BL-6J mice (Jackson Laboratories; ∼10 weeks old) were administered with 6 mg/ kg doxorubicin (in sterile saline solution; 0.9% *w/v* NaCl), intraperitoneally, once weekly for 3 weeks (cumulative dose of 18 mg/ kg). Animals were monitored during the administration and for a further 5 weeks with transthoracic echocardiographic measurements (Vevo® 2100 system, as detailed below) taken at baseline and weeks 1, 2, 3, 5 and 8. For ET treated animals, 70 mg/kg ET (dissolved in sterile saline; dose was established previously in [28] to significantly elevate levels in the heart) was administered by oral gavage daily for 1 week prior to administration of doxorubicin then once weekly for the 8-week duration of the study (Fig. 1A). Animals were assigned to 3 groups comprising saline control (no doxorubicin; n=7), doxorubicin alone (cardiotoxicity model; n=10) and doxorubicin with ET administration (evaluation of ET treatment; n=10). Mice were euthanized within 48h after the final echocardiogram and blood and tissues (heart, liver, kidney, and bone marrow) were collected for ET and biomarker analysis. In addition to measurements of cardiac function, the levels of ET and related metabolites, OCTN1 expression, and oxidative biomarkers were also measured in the blood and tissues of the animals.

**Figure 1:**
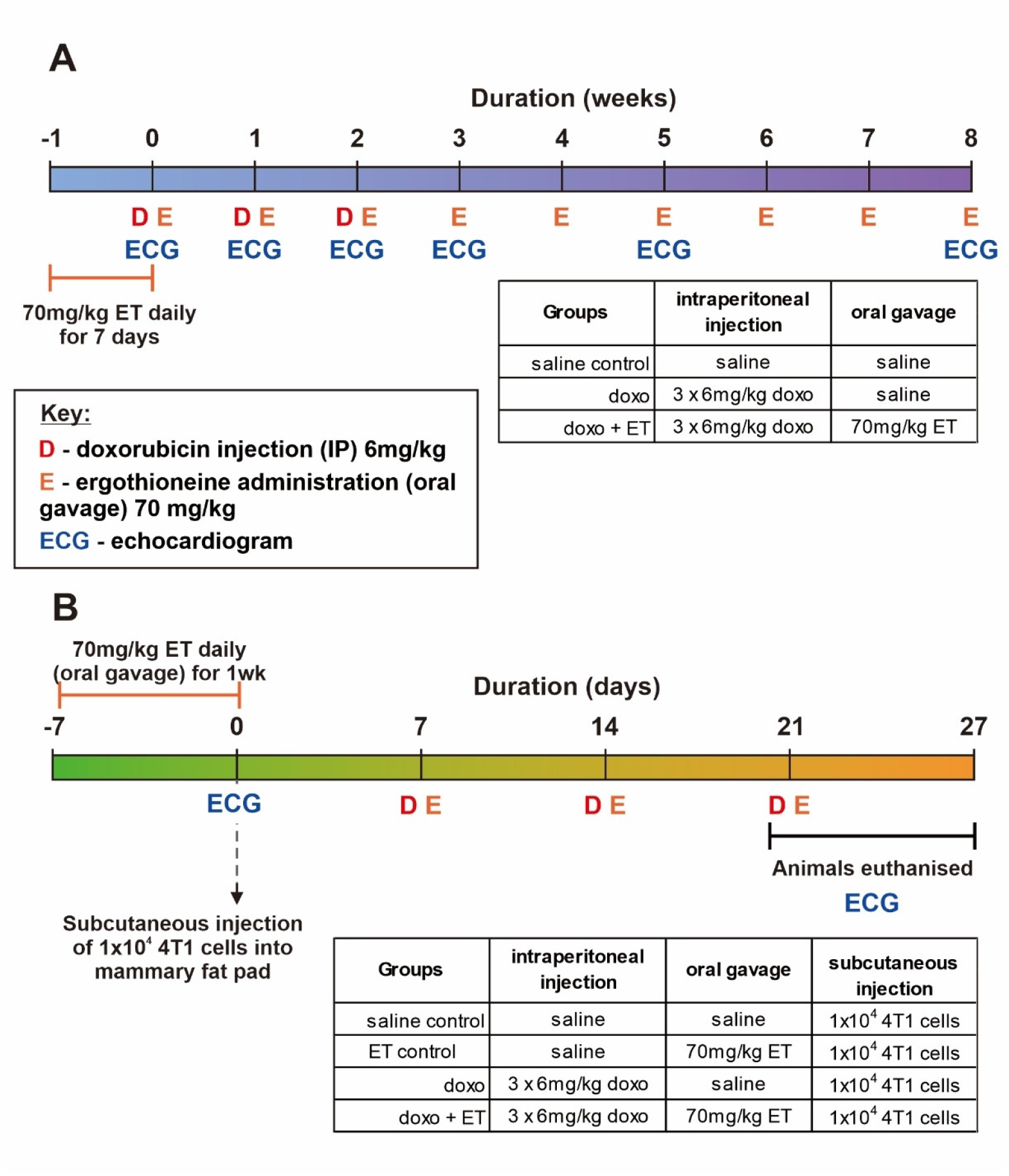
Timeline of animal studies and groups. (A) Doxorubicin-cardiotoxicity animal study in C57BL6 mice with or without ET administration. Mice (except for saline controls) were given 6 mg/kg doxorubicin weekly for 3 weeks and monitored periodically for a further 5 weeks. (B) A breast cancer model using injection of 4T1 cells into the mammary fat pad of BALB-C mice was used to evaluate the direct effect of ET on tumour growth and doxorubicin chemotherapy. Animals administered with doxorubicin were given 6mg/kg doxorubicin weekly for 3 weeks. In both studies, animals supplemented with ET were given 70 mg/kg ET daily for 7 days prior to the commencement of the study, then once weekly thereafter until the completion of the study.

### Mouse breast cancer model

A mouse breast cancer model was used to evaluate if ET interferes with the chemotherapeutic effect of doxorubicin and/or stimulates the growth of the cancer. BALB-C mice at 8 weeks of age were subcutaneously injected with 50 µl of a 4T1 cell (murine mammary gland carcinoma) suspension (containing 1×10^4^ cells) into the female mouse mammary fat pad. All mice were given the breast tumour injections and randomly assigned to groups (n=6/ group) given either saline (vehicle control), ET alone, doxorubicin alone, or doxorubicin with ET administration (Fig. 1B). For ET administration, 70mg/ kg were given to animals daily for 7 days prior to commencement of the study and once weekly until animals were euthanized. Animals administered doxorubicin were given 3 doses of 6mg/ kg intraperitoneally once weekly for 3 weeks, with the first dose commencing 1 week following injection of the breast cancer (Fig. 1B).

Only baseline and endpoint echocardiographic measurements were taken in this study due to the premature euthanasia of animals, to prevent undue stress to animals due to high dose of doxorubicin (required to suppress the aggressive tumour growth) or overgrown or ulcerated tumour in saline controls. However, these were sufficient to establish the cardiotoxic effects of doxorubicin with and without ET.

Mice were euthanized within 24h after the final echocardiogram and blood and tissues (heart, liver, kidney, and bone marrow) were collected for ET and biomarker analysis. The tumour size was measured with a vernier calliper after excision and subsequently weighed.

### Echo measurement and assessment

Animal cardiac function was assessed using a high frequency ultrasound system Vevo® 2100 (VisualSonics, Toronto, Canada) and analysed with Vevo® 2100 software, version 1.7.0, as established previously [50]. In brief, echocardiography was performed on mice under general anaesthesia (1-1.5% *v/v* isoflurane, Baxter, Singapore), cardiac left ventricular volumes and left ventricular ejection fraction (LVEF) were determined. To assess global cardiac dysfunction (both systolic and diastolic) in doxorubicin-induced cardiomyopathy, myocardial performance index (MPI), a parameter defined as the sum of the isovolumic contraction time and relaxation time divided by the ejection time, was determined by Doppler echocardiography of left ventricular inflow and outflow [51, 52]. Echocardiography was performed and analysed by a researcher blinded for the treatment of animals.

### Quantitative PCR of mouse heart RNA

RNA was isolated from heart samples using the NucleoSpin TriPrep kit with on-column DNase digestion (Macherey-Nagel, Dueren, Germany). Extracted RNAs were quantified and checked by 260/280nm measurement using the Synergy H1 spectrophotometer with Take 3 microvolume plate (BioTek; Agilent Technologies, CA, USA). Reverse-transcription PCR (RT-PCR) was performed using QuantiTect RT kit (Qiagen, Hilden, Germany), and quantitative PCR (qPCR) was performed using BlitzAmp Hotstart qPCR mix (MiRXES, Singapore) with an Applied Biosystem 7500 Real-Time PCR system (Thermo Fisher Scientific, MA, USA). Primers sequences are listed in Supplementary Data (Fig. S1), with gene expression normalised to β-actin.

### Extraction of blood and tissues ET and its metabolites

10µl of plasma, or ∼10mg of heart tissue (accurately weighed) were mixed with ET-d_9_ and hercynine-d_9_ internal standards and ultrapure water (Arium; Sartorius, Gottingen, Germany) to 100µl final volume. For plasma ET, samples were precipitated with 50µl of methanol at - 20°C overnight. For heart, tissue was homogenised with a motorized pellet pestle in 150µl of ultrapure water containing internal standards. After homogenization, 750µl of ice-cold methanol was added and samples were vortexed for 30s before incubating at −20°C overnight. Precipitated plasma and heart tissue samples were then centrifuged (20,000*g*, 15min, 4°C) and supernatants were evaporated under a stream of N_2_ gas before reconstituting in 100µl ultrapure water. Any debris was removed by centrifugation before transferring into silanised glass inserts with vials (Agilent CrossLab, CA, USA) for analysis by liquid chromatography mass spectrometry (LC-MS/MS). ET or hercynine levels were normalised against the accurate mass of tissue and expressed in mass per wet weight of tissue ± SEM.

### Quantifying biomarkers of oxidative damage to RNA and DNA

Levels of 8-hydroxyguanosine (8OHG) and 8-hydroxydeoxyguanosine (8OHdG), were measured in the heart tissue as biomarkers of oxidative damage to RNA and DNA, respectively. Approximately 20mg of heart tissue was used to extract RNA and DNA using TRIzol reagent (Invitrogen, MA, USA) as per the manufacturer’s protocol. All procedures were performed on ice to reduce oxidation artefacts. Isolated RNA was dissolved in RNase-free water, while DNA was dissolved in 160µl of 10mM Tris pH 8.0 buffer at 4°C overnight. 50µg of RNA were enzymatically hydrolysed at 37°C using 20µg RNase A, 1mU phosphodiesterase (P3242; Sigma-Aldrich, MO, USA), and 2U alkaline phosphatase (A2356; Sigma-Aldrich, MO, USA) in buffer (10mM Tris pH 8.0, 5mM MgCl2) in a volume of 100µl for 1h. Isolated DNA was hydrolysed at 37°C with 1U of benzonase (Merck, NJ, USA), 3mU phosphodiesterase, and 4U alkaline phosphatase in 200µl of buffer for 6h. Internal standards were added before hydrolysis. The reaction was quenched with 5 volumes of ice-cold methanol, vortexed, and stored in −20°C overnight. All samples were centrifuged (20,000*g*, 15min, 4°C) and supernatants were evaporated under a stream of N_2_ gas before reconstituting in 60µl of ultrapure water. Any precipitates were removed by centrifugation before transferring into silanised glass inserts with vials for LC-MS/MS analysis.

### Protein carbonylation ELISA

Diluted plasma (25x in PBS) and TRIzol extracted heart proteins were derivatised with 4mM DNPH (in 2M HCl) as per manufacturer’s instructions; Oxyblot Protein Oxidation Detection Kit (Merck Millipore, MA, USA). 2µg of proteins in 200µl 100mM pH 9.6 bicarbonate buffer were left to bind to the ELISA plate overnight at 4°C. The plate was washed 5 times with PBS and blocked with 100µl Pierce Protein-Free T20 buffer for 1h. Plate was washed with PBS-T (0.1% Tween-20) and incubated with 100µl 1:200 primary antibody (Oxyblot kit) for 2h with shaking. Plate was then washed and incubated with 100µl 1:400 secondary antibody (Oxyblot kit) for 1h. Plate was washed and 100µl of TMB (Abcam, Cambridge, UK) was added. The reaction was stopped with 100µl of 2M H_2_SO_4_ and absorbance determined at 450nm using the Synergy H1 spectrophotometer (BioTek; Agilent Technologies, CA, USA). Carbonylation levels were quantified against a standard curve.

### Liquid chromatography mass spectrometry analysis

LC-MS/MS was carried out using an Agilent 1290 UPLC system coupled to an Agilent 6460 triple quadrapole mass spectrometer (Agilent Technologies, CA, USA). Samples were kept at 10°C in the autosampler during analysis.

For ET and hercynine analysis, 2µl of the processed samples were injected into a Cogent Diamond-Hydride column (4µm, 150 × 2.1mm, 100Å; MicroSolv Technology Corporation, NC, USA) maintained at 40°C. Solvent A was acetonitrile in 0.1% *v/v* formic acid, and Solvent B was 0.1% *v/v* formic acid in ultrapure water. Chromatography was carried out at a flow rate of 0.5ml/ min using the following gradient elution; 1min of 20% solvent B, following by a gradual increase to 40% solvent B over 3min to elute ET. Solvent B was further increased to 90% over 1min to elute hercynine, and this was maintained for 3.5min before returning to 20% for 3.5min to re-equilibrate the column. The total run time was 12min. The retention times for ET and hercynine were 4.2 and 6.8 min, respectively.

For the 8OHG and 8OHdG analysis, 10 µl of the processed samples were injected into an Accucore PFP column (150 × 3.0mm, 2.6µm; Thermo Fisher Scientific, MA, USA) maintained at 30°C. Solvent A was acetonitrile, and Solvent B was 0.1% *v/v* formic acid in ultrapure water. Chromatography was carried out at a flow rate of 0.5 ml/min using the following gradient elution; 1.5min of 98% solvent B, followed by a 4min gradual decrease of solvent B to 95%. The column was washed with 5% solvent B for 2.5min and re-equilibrated with 98% solvent B for 4min. The total run time was 12min. The retention times for guanosine, 8OHG, dG, 8OHdG were 3.7, 4.3, 4.8 and 6.1min, respectively.

For all targets, MS was carried out under positive ion ESI and multiple reaction monitoring mode. Capillary voltage was set at 3200V, and gas temperature was kept at 350°C. Nitrogen sheath gas pressure for nebulizing sample was at 50psi, and gas flow set at 12L/min. Ultra-high purity nitrogen was used as collision gas. The optimised precursor to product ion transitions and their respective fragmentor voltages and collision energies are listed in Supplementary Data (Fig. S2).

### Statistical analysis

Data were tabulated using Microsoft Excel (Microsoft Corporation, WA, USA). Statistical analyses (1-way analysis of variance (ANOVA) with Tukey’s multiple comparison test or 2-way ANOVA, with multiple column comparison) were performed, and graphs were generated using GraphPad Prism version 9.2.0 (GraphPad Software, CA, USA). Unless otherwise stated, all data are expressed as mean ± standard error, with *P* < 0.05 considered statistically significant.

## Results

### Animal weights in doxorubicin treated mice

Administration of 6 mg/kg doxorubicin weekly for 3 weeks (total cumulative dose of 18 mg/kg) in C57BL-6J mice led to significant decreases in animal weights (Fig. 2) compared to saline treated controls whose weights continued to increase over time. Animal weights in doxorubicin treated animals remained low for the entire duration of the 8-week study, even after doxorubicin administration had ceased. Doxorubicin treated animals also supplemented with 70mg/kg ET also declined in weight however their weights were able to gradually recover after doxorubicin administration had ceased, which was significantly greater than doxorubicin alone by week 8.

**Figure 2:**
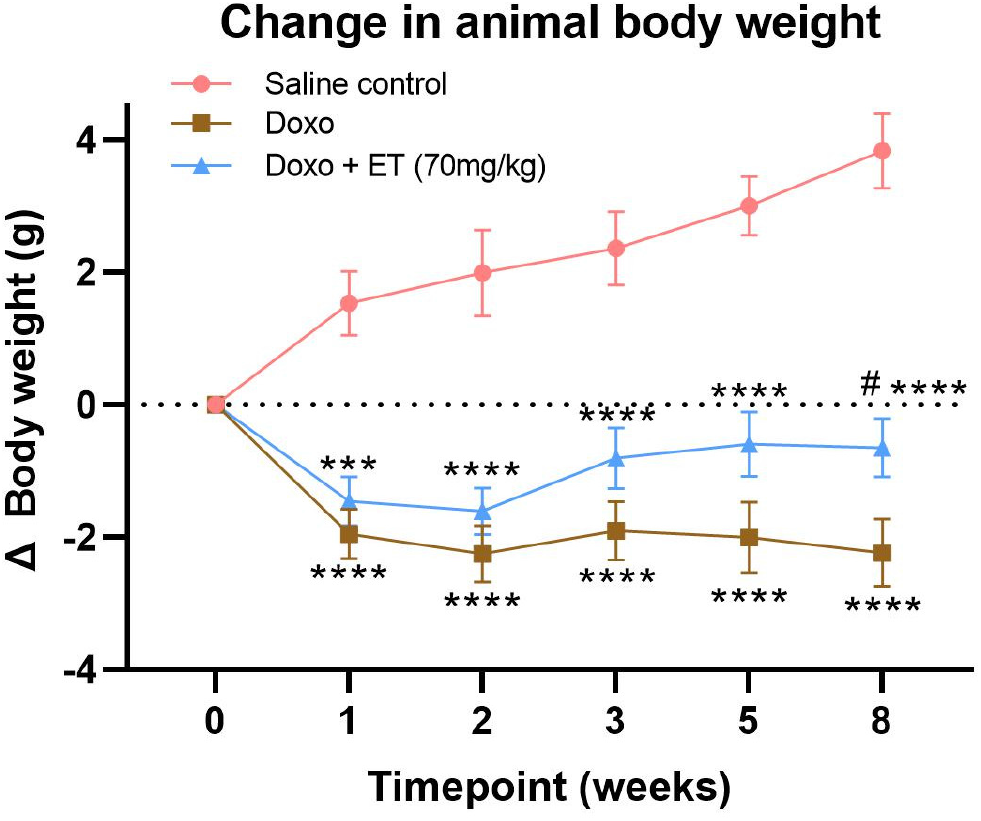
Animal weights. Animal weights were taken at each timepoint to compare weights of animals with doxorubicin administration with (doxo + ET) and without (doxo) ET supplementation with untreated (saline control) animals. Mice administered with doxorubicin had a significantly decline in mean body weight compared to saline controls. While doxorubicin treated animals supplemented with ET also had significant declines in body weight their body weights were able to recover being significantly greater than doxorubicin treated animals alone by week 8. 2-way analysis of variance (ANOVA) using multiple column comparison with significance **** *P* < 0.0001, *** *P* < 0.001 vs. saline control and # *P* < 0.05 vs. doxorubicin alone.

### Echocardiographic assessment of cardiac function in doxorubicin treated mice

Left ventricular systolic function as assessed by echocardiographic assessment of the change in LVEF is a primary measure of cardiac function. As expected, the LVEF in saline treated controls remained constant relative to baseline over the 8-week study (Fig. 3A). In contrast, administration of doxorubicin significantly decreased LVEF from baseline (ΔEF) from second week onward (corresponding to second dose of doxorubicin; Fig. 3A). To comprehensively evaluate the impact of doxorubicin on cardiac function, we also performed cardiac Doppler echocardiography to determine MPI that reflexes myocardial performance in both systolic and diastolic aspects [51, 52]. As shown in Fig. 3B, a significant increase in MPI was observed in doxorubicin treated mice relative to saline treated controls, indicating impaired cardiac function by administration of doxorubicin in mice. Remarkably, the oral supplementation of mice with 70mg/kg ET prior to doxorubicin administration, as illustrated in Fig. 1A, prevented doxorubicin-induced cardiac dysfunction, as evidenced by preservation of LVEF and a significantly lower increase in MPI (Fig. 3A & B).

**Figure 3:**
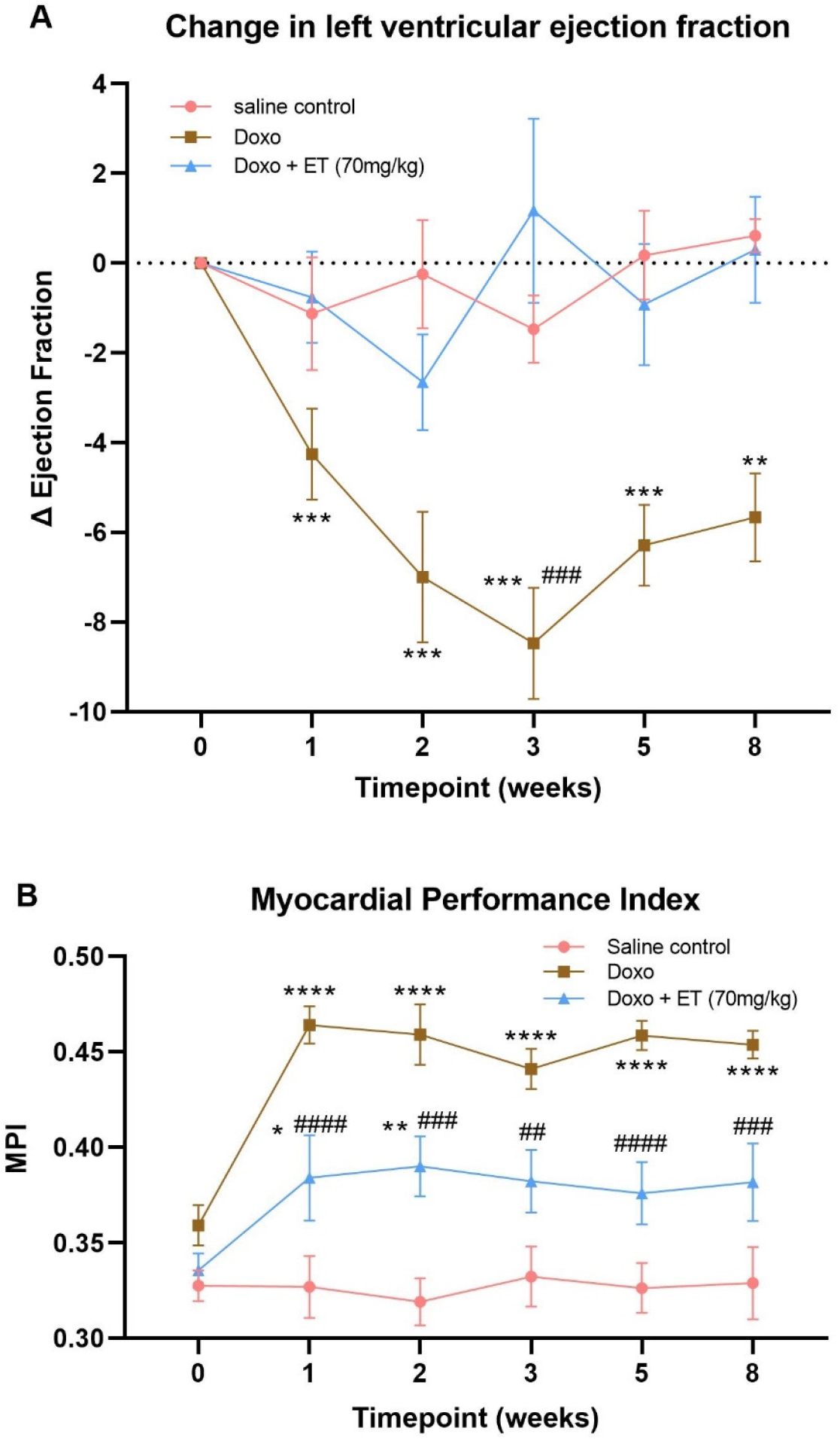
Assessment of cardiac function. (A) The change in LVEF was measured as an indicator of cardiac (systolic) function. Significant decreases in LVEF were seen with doxorubicin (alone) administered animals which was significant versus control from 2 weeks. Supplementation of doxorubicin treated animals with ET prevented this decline in LVEF which was significantly different to doxorubicin treated alone at the 3-week time point. (B) Similarly myocardial performance index was significantly increase in doxorubicin treated animals (indicating declining cardiac function). ET administration decreased MPI but not to baseline/ saline control levels. 2-way ANOVA using multiple column comparison with significance **** *P* < 0.0001, *** *P* < 0.001, ** *P* < 0.01, * *P* < 0.05 vs. saline control and # *P* < 0.05 vs. doxorubicin alone.

### Expression of ET transporter mRNA in heart

A trend to an increase in mRNA expression of OCTN1 was observed in the heart following administration of doxorubicin compared with saline controls, however this was not significant (Fig. 4A). No change was observed in OCTN1 expression in animals supplemented with ET prior to administration of doxorubicin, when compared to saline controls.

**Figure 4:**
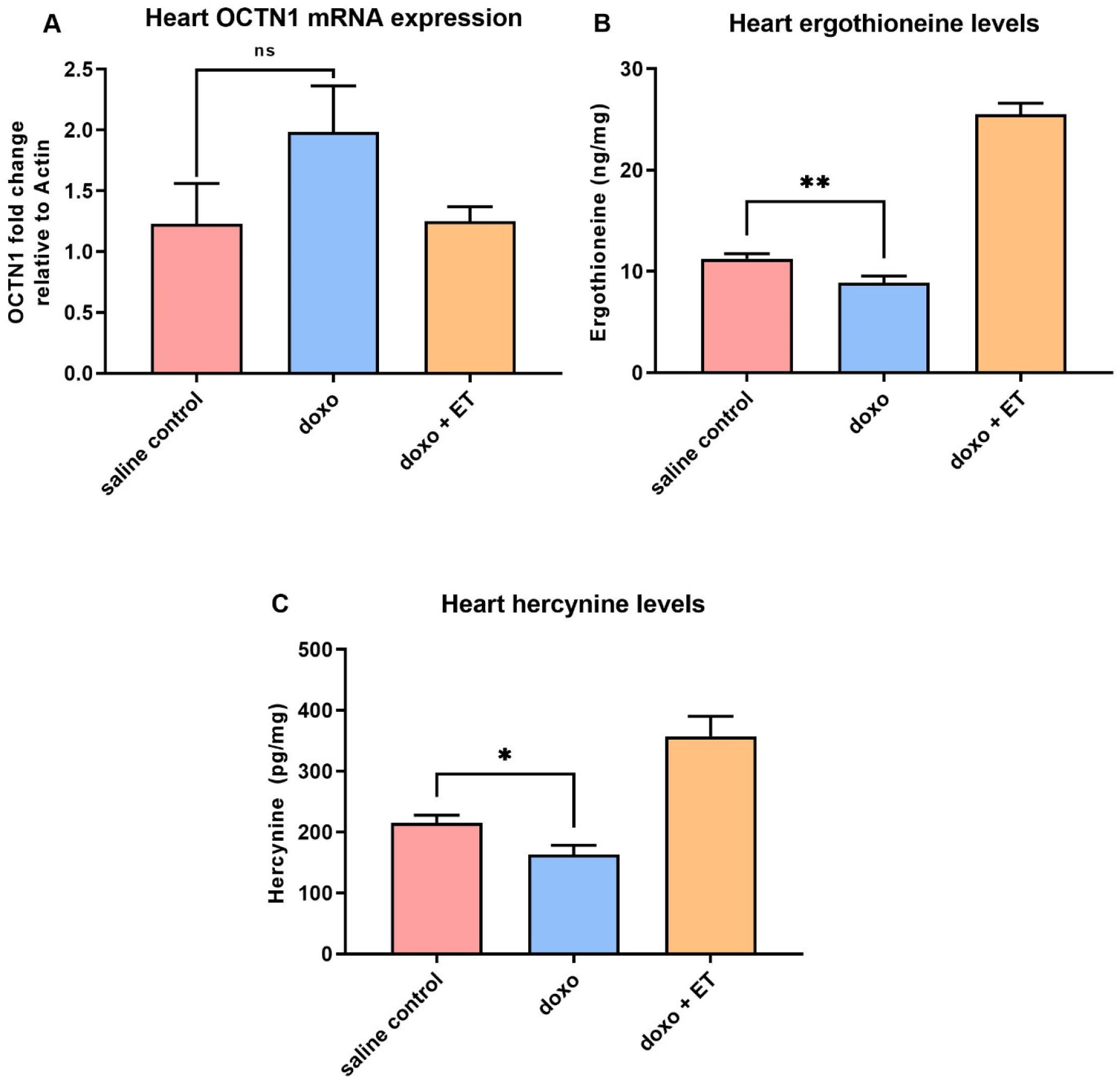
Levels of ergothioneine and hercynine in the heart. (A) Real-time PCR analysis of OCTN1 (the ET transporter) mRNA expression in heart tissue. Although a slight increase in OCTN1 mRNA was seen with doxorubicin administration, this was not significant. No changes were seen in doxorubicin treated animals with ET supplementation. (B) Significantly lower levels of ET were seen in the heart with doxorubicin treatment. This may be due to increased consumption of ET as it scavenges ROS. The heart levels of hercynine (C) also decrease significantly relative to saline control. Supplementation with ET as expected increases levels of hercynine (prior studies show that levels of hercynine correlated closely with ET [59]). 1-way ANOVA with Tukey’s multiple comparison test; ** *P* < 0.01, * *P* < 0.05 vs. saline control; *P* < 0.0001 saline control or doxorubicin alone vs. doxorubicin + ET groups in (B) & (C); ns – not significant.

### Levels of ergothioneine and hercynine in heart tissues

It should be noted that as with prior studies (Tang et al. [28]), detectable basal levels of ET are present in both blood and heart due to the avid accumulation and retention from the trace levels present in the standard mouse diet. Doxorubicin administration significantly decreased levels of ET in the heart (Fig. 4B), which is presumably through utilization (i.e. oxidation). Levels of the ET metabolite hercynine also decreased with doxorubicin administration alone (Fig 4C). As expected, oral supplementation of ET, significantly increased the levels of ET (∼2.5 fold higher than baseline; Fig. 4B) as well as levels of hercynine (Fig. 4C).

### Levels of ergothioneine and hercynine in plasma

The levels of ET in the plasma also follow a similar trend to the heart tissues, with a significant drop in ET levels following doxorubicin administration (Fig. 5A). However, levels of hercynine (a metabolite of ET) had a trend to increase, although this was not significant, which could be due to efflux from tissues (Fig. 5B). Again, supplementation with ET significantly raised the levels of ET (∼3-fold increase) as well as hercynine in the plasma.

**Figure 5:**
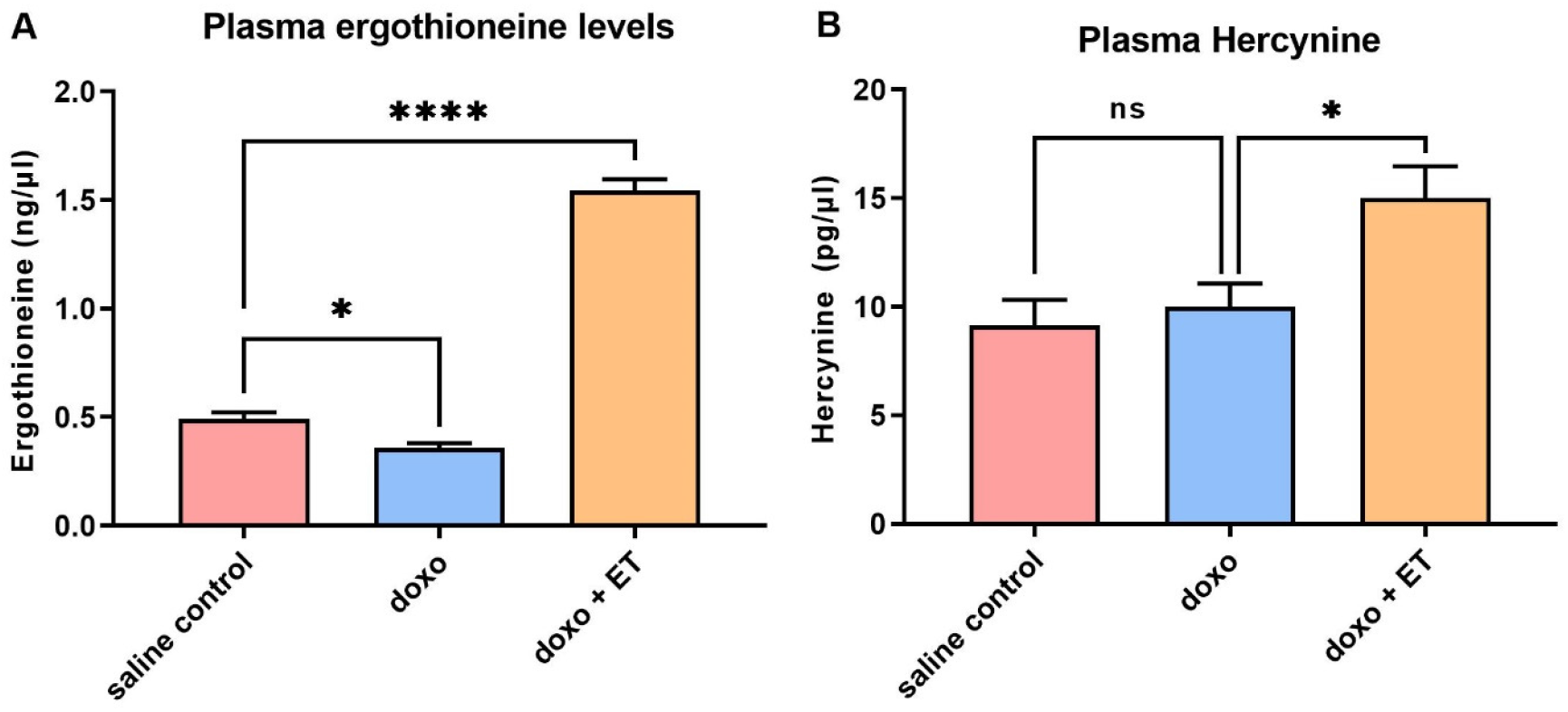
Plasma ergothioneine and hercynine. A significant decrease was seen in plasma (A) ET levels with administration of doxorubicin relative to saline controls, similar to what was observed in the heart. A non-significant trend to increase in plasma hercynine was observed following doxorubicin administration (B). As expected, supplementation increased plasma levels of ET and hercynine. 1-way ANOVA with Tukey’s multiple comparison test; (A) * *P* < 0.05, **** *P* < 0.0001 vs. saline controls; (B) * *P* < 0.05 vs doxorubicin alone.

### Oxidative damage biomarkers

The levels of 8OHdG, 8OHG, and protein carbonyls were measured in the heart tissues as established biomarkers of oxidative damage to DNA, RNA and protein, respectively [11, 53]. A slight but non-significant increase in DNA and RNA oxidation was observed with doxorubicin administration (Fig. 6A & B), while supplementing ET appeared to decrease DNA and RNA oxidation markers but again this was not significant relative to animals treated with doxorubicin alone. No apparent trends were observed for protein carbonyls in the heart tissue (Fig. 6C); however, plasma protein carbonyls appear to follow a similar non-significant trend as DNA and RNA oxidation markers in the heart (Fig. 6D).

**Figure 6:**
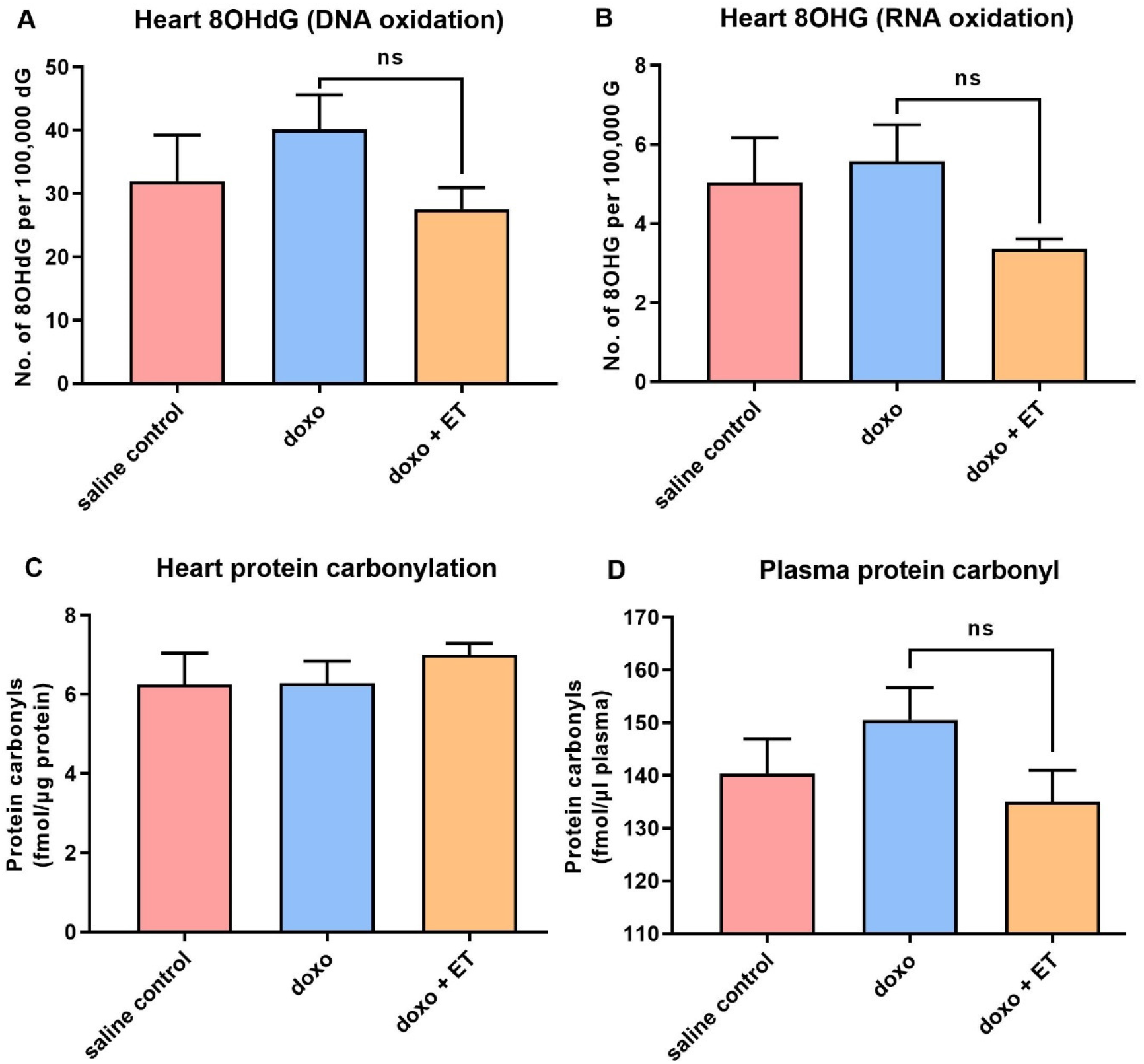
Oxidative damage biomarkers. Although slight increases were observed in the quantification of 8OHdG, a biomarker of oxidative DNA damage (A), and 8OHG, a biomarker of oxidative RNA damage (B), with doxorubicin administration relative to saline controls, this was not significant (ns; 1-way ANOVA, Tukey’s multiple comparison test). Supplementation with ET also appeared to decrease oxidative markers of both DNA and RNA, but this was not significant (A&B). No differences were seen in heart protein carbonyls (C) between any of the groups. A similar trend in plasma protein carbonyls (D) was seen as with nucleic acid oxidation markers in heart but again this was not significantly different.

### Breast cancer model

In BALB-C breast cancer model, the body weights of animals between saline and ET control groups (without doxorubicin) were not different. However, doxorubicin administration caused a significant loss of body weight and premature death in these animals, regardless of ET supplementation (Fig. 7A). As with the C57BL-6J doxorubicin cardiotoxicity model, the administration of doxorubicin to the BALB-C mice led to a significant decrease in LVEF relative to baseline (Fig. 7B). Supplementation with ET almost completely abolished this doxorubicin-induced decrease in LVEF, and animals fared significantly better than animals with doxorubicin administration alone. These results indicate that ET protects against doxorubicin-induced cardiac dysfunction even in the presence of breast cancer.

**Figure 7:**
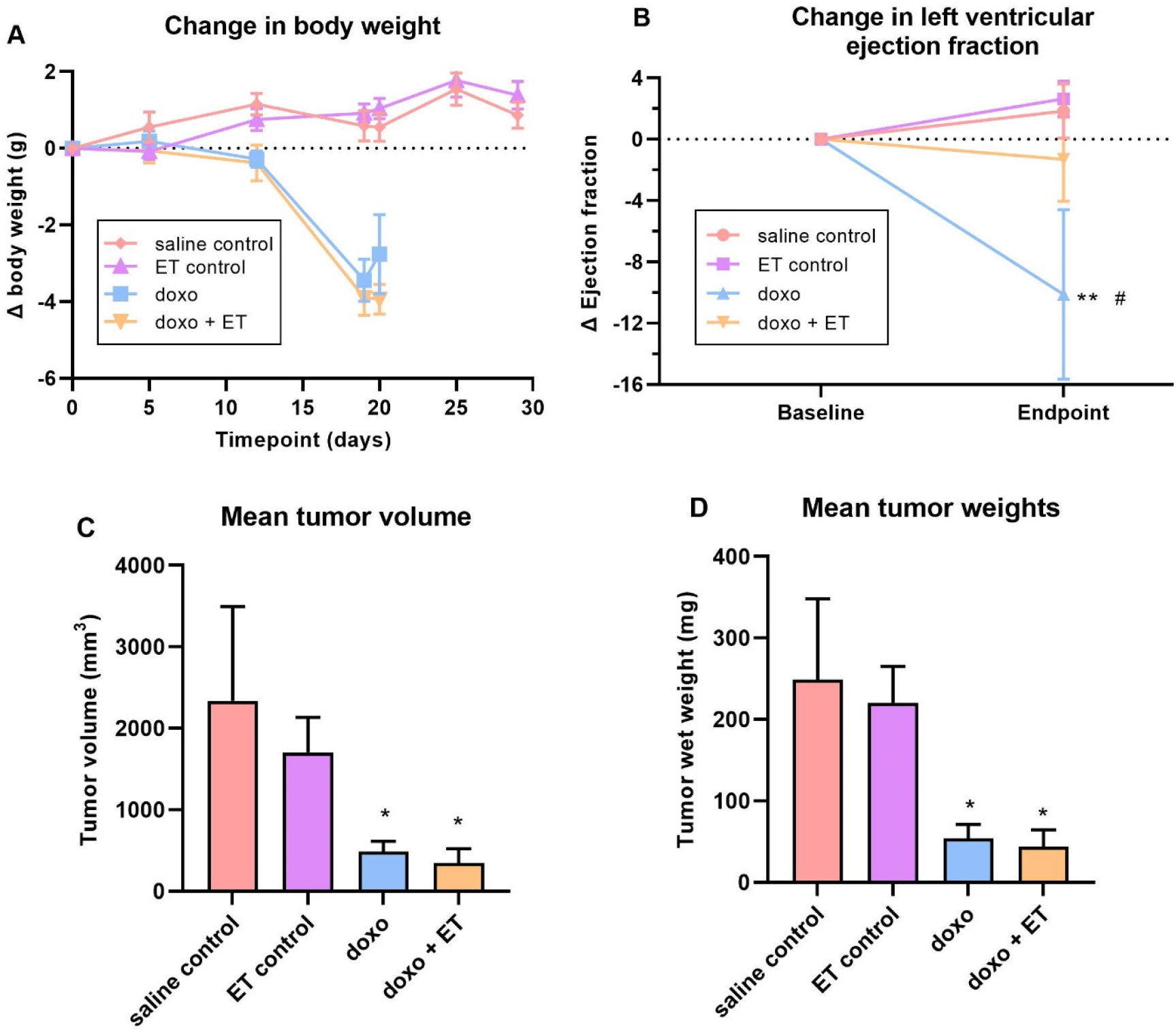
Breast cancer model. (A) Administration of doxorubicin resulted in a significant (*P <* 0.0001; 1-way ANOVA multiple comparison test) decline in body weights with or without ET supplementation relative to saline controls. ET supplemented animal weights remained akin to saline controls. (B) Cardiac function (LVEF) was measured at baseline and endpoint in the breast cancer model. As with the C57BL-6J doxorubicin model, a significant decrease in LVEF was seen with administration of doxorubicin relative to saline control animals. Supplementation with ET however significantly prevented this decline in cardiac function (2-way ANOVA with multiple comparison test; ** *P* < 0.01 vs. saline control # *P* < 0.05 vs. doxo + ET). The average volume (C; estimated based on calliper measurements) and weight (D) of the excised tumours are shown. Saline control animals had the largest tumour size, but administration of doxorubicin significantly decreased volume/weight of the tumour (* *P <* 0.05; 1-way ANOVA with multiple comparison test). Supplementation of animals with ET alone had a slight, but not significant, decrease in tumour size compared to saline controls. Likewise, ET supplementation in doxorubicin treated animals had a slight but non-significant decrease in tumour size relative to doxorubicin alone. This indicates that ET does not aggravate tumour growth nor interfere with the chemotherapeutic effect of doxorubicin.

Following euthanasia, the excised breast tumours were measured with a calliper to estimate the volume and subsequently weighed. The estimated volume (Fig. 7C) and mass of the tumours (Fig. 7D) were closely correlated. The tumour size and mass in saline control animals were the greatest, but doxorubicin administration significantly reduced the volume and mass of the tumour. Interestingly a slight decline in tumour size was observed in animals supplemented with ET alone or with doxorubicin when compared with saline controls or doxorubicin alone, respectively, however this was not significant (Fig. 7C & D). Supplementation of doxorubicin treated animals with ET thus did not appear to affect the chemotherapeutic efficacy of doxorubicin.

## Discussion

Cardiotoxicity is one of the leading causes of death in patients undergoing or having undergone cancer treatment [54]. Indeed, anthracyclines which are commonly applied as chemotherapy drugs against leukaemia and breast cancer (although they also find application in many soft tissue cancers and tumours of the stomach, lung, ovaries, and other organs), are known to be associated with cumulative dose-related cardiotoxicity (i.e. risk of cardiotoxicity increases with total cumulative dose). Anthracycline-induced cardiomyopathy is mostly irreversible and may lead to congestive heart failure. In breast cancer survivors, cardiovascular disease accounts for 35% of non-cancer related deaths in survivors aged ≥50 years [55]. Although some cases of acute cardiotoxicity, occurring during or soon after anthracycline administration, most cases are chronic cardiotoxicity occurring within the first year (early-onset) or decades following completion of chemotherapy (late-onset) [56]. Up to 65% of adult survivors of late-onset cardiotoxicity, particularly relevant to treatment of childhood malignancies with doxorubicin, present evidence of left ventricular contractile abnormalities [57]. Furthermore, concomitant application of certain other anti-cancer drugs (despite absence of cardiotoxicity in those drugs alone) may further exacerbate anthracycline cardiotoxicity and increase the risk of cardiac dysfunction [58].

We established that an optimised cumulative dose of 18 mg/ kg doxorubicin over 3 weeks was able to induce cardiotoxicity and significantly decrease LVEF in the C57BL-6J mice (Fig. 3A). Although the EF partially recovered over the 5 week follow up period this remained significantly lower than control mice for the entire study. Supplementing this doxorubicin cardiotoxic mouse model with ET (70 mg/kg; daily for 7 days prior to doxorubicin administration and subsequently once weekly *via* oral gavage) abolished the doxorubicin-induced decrease in LVEF. This dose was previously established in mice [28] demonstrating differential distribution to tissues of the animal including the heart. By endpoint a ∼2.5-fold increase in ET levels was observed in the heart tissue compared with saline controls, with ET supplementation (Fig. 5A). This is a critical point as many antioxidants or therapeutics may not possess a specific transporter or may be rapidly metabolised and/or excreted leading to poor bioavailability. We have similarly observed the accumulation and strong retention of ET in human subjects following oral supplementation [59], with transcriptomics revealing the presence of *slc22a4* (gene encoding OCTN1) mRNA in human heart [25], suggesting a similar accumulation of ET as with mice. Indeed, the uptake of ET is dictated by the expression of OCTN1 and knocking out this transporter in mice leads to tissues devoid of ET [60], which seems to predispose tissues to oxidative stress and inflammation [61-63]. Conversely, we hypothesised that tissue stress and injury may feedback to upregulate OCTN1 expression and increase ET uptake at the site of tissue injury [48, 49]. Supporting this, a metabolomic study revealed a significant increase in ET levels in the heart following pressure overload (by aortic constriction) or myocardial infarction [47]. As seen in our cardiotoxicity model, doxorubicin administration alone tended to increase the expression of OCTN1 mRNA, although this was not significant (Fig. 4A). However, when high levels of ET were present (due to oral supplementation), the induction of OCTN1 expression was not observed. As the tissues were collected at endpoint, it is possible that the expression levels of OCTN1 had already declined since this was more than 5 weeks following the final doxorubicin administration. This stabilisation is evident from observing the animal weights which significantly decline during doxorubicin administration with or without ET (Fig. 2), but somewhat recover following cessation of doxorubicin administration. Animals supplemented with ET appeared to recover faster (from 3-week timepoint onwards) and their mean body weights were significantly higher compared with doxorubicin treated animals alone.

Doxorubicin administration also significantly increased MPI scores, an assessment of global cardiac dysfunction which combines both systolic and diastolic performance [64], relative to saline controls (Fig. 3B). The supplementation with ET in doxorubicin treated animals significantly lowered this doxorubicin-induced increase in MPI score. However, ET supplementation did not reduce this to levels of the saline controls possibly indicating a subclinical left ventricular cardiotoxicity [65]. Perhaps prolonged ET administration or longer observation period is required to demonstrate the efficacy of ET against late-onset cardiac dysfunction (which can occur decades after completion of anthracycline chemotherapy in humans).

A significant decrease in basal heart levels of ET (compared to saline controls) was observed with doxorubicin administration alone, despite the trend to an increase in OCTN1 expression observed (Fig. 4A). This may be indicative of ET utilisation/ oxidation in the heart. Indeed, doxorubicin-mediated iron overload in mitochondria and mitochondrial redox cycling have been shown to produce excessive ROS [5, 11]. ET has been previously shown to scavenge a range of radicals including singlet oxygen [66], hydroxyl radicals [31], peroxynitrite [67], or ferryl complexes [46, 68], and has also been shown to protect against damage by hydrogen peroxide [69, 70] and superoxide [71, 72]. Hercynine is the purported stable end-product of ET oxidation (*via* ergothioneine sulphonate [73]). Indeed, we have shown animal tissues devoid of ET in OCTN1 knockout mice also lack hercynine, indicating it is a metabolite of ET [60]. Levels of hercynine in the heart tissues were seen to decline significantly following doxorubicin administration alone and significantly increase following ET administration (Fig. 4C) in a similar trend to ET levels in the heart. This is not surprising given that we have previously shown that hercynine levels closely correlate to levels of ET [59]. In contrast to ET, hercynine is unlikely to be retained in the cardiomyocytes and certainly any transient increase in hercynine due to ET oxidation may not be detectable by the endpoint (5-weeks post doxorubicin) when the tissues were collected. Supporting this the levels in plasma appear to be slightly elevated (Fig. 5B), although this is not significant, possibly due to excretion from the body. Another possible explanation is that hercynine may not be the final oxidation product, and may react with oxidants such as hypochlorite as was suggested recently [74].

A trend of increasing oxidative damage to DNA and RNA (Fig. 6A & B) was observed with administration of doxorubicin, while supplementation of doxorubicin treated animals with ET appeared to decrease levels of DNA and RNA oxidative markers, however in both cases this was not significant. Unfortunately, as the tissues were only collected at endpoint (8 weeks) to allow for observation of cardiac function following the cessation of doxorubicin administration, this meant that earlier increases in oxidative damage biomarkers may have normalised. With highly active cellular DNA repair mechanisms [11] and high RNA and protein turnover it is possible that the peak of oxidative injury was not captured. As such it is difficult to ascertain if ET is protecting the heart due to a decrease in DNA, RNA or protein oxidative damage either mediated through chelation of Fe^2+^ in cardiomyocytes or prevention of mitochondrial dysfunction. Iron plays a key role in doxorubicin cardiotoxicity, through the formation of doxorubicin-Fe complexes [75] and iron-overload in mitochondria. Animal studies found that excessive iron exacerbates doxorubicin cardiotoxicity [76, 77]. Conversely, iron chelators such as dexrazoxane and deferoxamine, have been shown to decrease cardiotoxicity in animal models and some clinical studies [76]. ET can also bind divalent metal ions such as Fe^2+^and Cu^2+^ [32], forming redox-inactive complexes preventing damage to critical cellular components such as DNA and mitochondria [33]. Indeed, the protective role of ET during ischemia-reperfusion is suggested to be in part mediated by iron chelation [78]. Similarly, the chelation of iron could prevent the association with doxorubicin and formation of complexes and/or inhibit its ability for partake in redox activities. However, further studies are needed to evaluate the exact cardioprotective mechanism(s) of ET.

In our breast cancer model, supplementation with ET did not have any significant effect on tumour size or weight, both alone or with doxorubicin, indicating ET does not exacerbate the tumour growth, nor does it interfere with the chemotherapeutic efficacy of doxorubicin, respectively, unlike other antioxidants [19-21]. This is especially important to establish. It is interesting to note that while not significant, ET administration itself slightly decreased the size of the tumour both in the absence and presence of doxorubicin. This suggests that ET alone may have a slight anti-tumour effect and may warrant further studies. Indeed, studies by D’Onofrio *et al*. [79], noted a dose-dependent anti-cancer effect of ET in colorectal cancer cells, by inducing necroptosis through SIRT3/MLKL pathways.

While further studies are needed to elucidate the mechanisms of protection by ET, this study paves the foundations for future clinical evaluation of ET as a cardioprotectant against anthracyclines in chemotherapy. Furthermore, ET is thought to be safe for clinical use, bestowed with the GRAS (generally recognized as safe) status by the US FDA and has received the European Food Safety Authority (EFSA) approval for use as a food supplement even in infants and children [80, 81].

## Conclusion

Anthracyclines such as doxorubicin remain a primary chemotherapeutic against haematological malignancies and solid tumors such as breast cancer, due to their effectiveness. However, counteracting the devastating cardiotoxic effects remains a major challenge. We demonstrated that the unique cytoprotective thione, ET, which can accumulate in heart tissues, decreases the cardiotoxic effects of doxorubicin in our mouse model. Moreover, using a murine breast cancer model we established that ET did not interfere with chemotherapeutic efficacy of doxorubicin nor exacerbate the growth of the breast tumour. Although further work is still needed to elucidate the protective mechanisms against doxorubicin, this study highlights the potential for ET to be used as a potential agent for alleviating anthracycline cardiotoxicity in cancer patients and certainly warrants further attention.

## Abbreviations

EFSA: European Food Safety Authority
ET: ergothioneine
EF: ejection fraction
FDA: (US) Food and Drug Administration
GSH: reduced glutathione
LC-MS/MS: liquid chromatography tandem mass spectrometry
LVEF: left ventricular ejection fraction
MS: mass spectrometry
OCTN1: organic cation transporter novel type-1
8OHdG: 8-hydroxydeoxyguanosine
8OHG: 8-hydroxyguanosine
SPE: solid phase extraction
ROS: reactive oxygen species

## Declaration

All authors declare no conflict of interest.

## Acknowledgements

The authors would like to thank the Ministry of Education AcRF Tier 1 (NUHSRO/2017/055/T1), the Tan Chin Tuan Centennial Foundation, the Ministry of Health – National Academy of Medicine Healthy Longevity Catalyst Award (HLCA20Jan-0057), and the National Medical Research Council (Individual Research Grant NMRC/1264/2010/082/12 and NMRC/OFTIRG/0081/2018) for providing financial support. Suet Yen Chong would like to thank the ESR/TENG GL PhD scholarship program for their support. The authors also wish to thank Tetrahedron (Paris, France) for the kindly providing the L-ergothioneine, L-ergothioneine-d_9_, L-hercynine, and hercynine-d_9_, used in our studies.

