## Supplementary Fig. S1 for "Protection against doxorubicin-induced cardiotoxicity by ergothioneine"

***Supplementary data***

**Figure S1:** Forward and reverse primer sequences used for qPCR.

| Genes | Forward Primer 5'→3' | Reverse Primer 5'→3' |
|---|---|---|
| **OCTN1** | AAGTCCTCTTTGCAACCATGGC | CTGTGTTGTTTGCCATTGTGGG |
| **β-ACTIN** | TGACAGGATGCAGAAGGAGA | CGCTCAGGAGGAGCAATG |

**Figure S2:** Mass spectrometry multiple reaction monitoring (MRM) transitions as well as the optimised fragmentor voltages and collision energies for those transitions.

| Targets | MRM transition | Fragmentor Voltage (V) | Collision Energy (eV) |
|---|---|---|---|
| ET | 230 → 186 | 103 | 9 |
| ET-$d_9$ | 239 → 195 | 98 | 9 |
| Hercynine | 198 → 95 | 94 | 21 |
| Hercynine-$d_9$ | 207 → 95 | 103 | 9 |
| 8OHdG | 284 → 168 | 80 | 10 |
| 8OHdG-$^{13}C_2$,$^{15}N_1$ | 287 → 171 | 80 | 10 |
| 8OHG | 300 → 168 | 85 | 10 |
| 8OHG-$^{13}C_1$,$^{15}N_2$ | 303 → 171 | 85 | 10 |
| dG | 268 → 152 | 70 | 6 |
| dG-$^{13}C_1$,$^{15}N_2$ | 271 → 155 | 65 | 5 |
| Guanosine | 284 → 152 | 70 | 10 |
| Guanosine-$^{15}N_5$ | 289 → 157 | 85 | 10 |

**Figure S3:** Image of excised tumours from each animal.

A

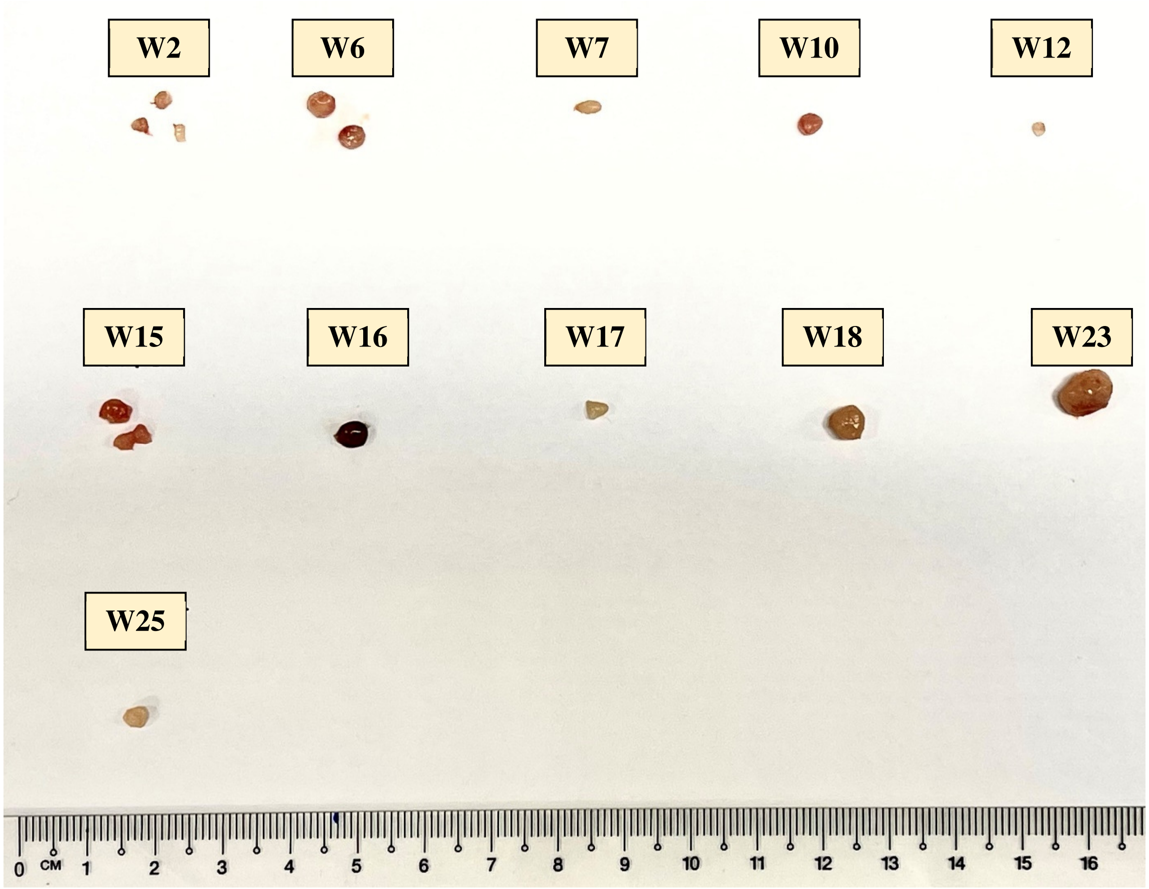


B

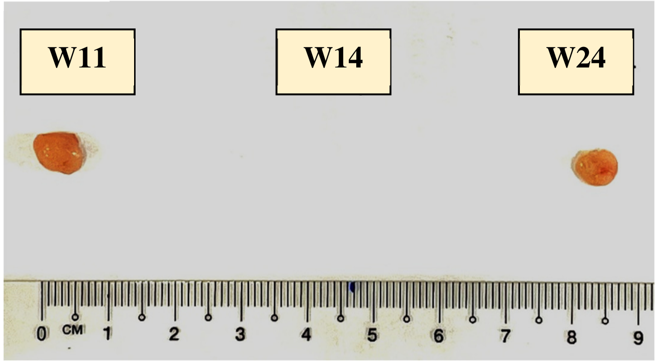

C

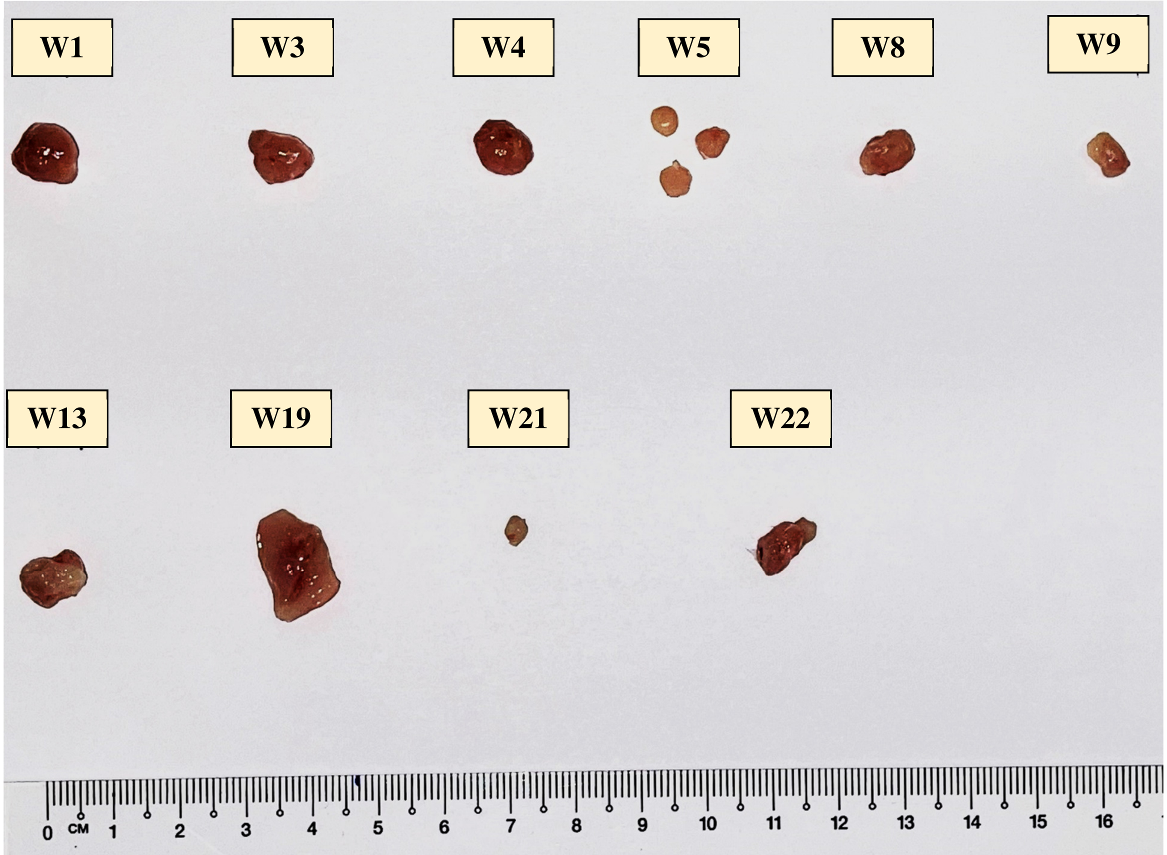


| Animal | Group | ET | doxorubicin |
|---|---|---|---|
| W1 | 2 | 70mg/kg | Saline |
| W2 | 4 | 70mg/kg | 6mg x 3 |
| W3 | 2 | 70mg/kg | Saline |
| W4 | 2 | 70mg/kg | Saline |
| W5 | 2 | 70mg/kg | Saline |
| W6 | 4 | 70mg/kg | 6mg x 3 |
| W7 | 4 | 70mg/kg | 6mg x 3 |
| W8 | 2 | 70mg/kg | Saline |
| W9 | 2 | 70mg/kg | Saline |
| W10 | 4 | 70mg/kg | 6mg x 3 |
| W11 | 4 | 70mg/kg | 6mg x 3 |
| W12 | 4 | 70mg/kg | 6mg x 3 |

| Animal | Group | ET | doxorubicin |
|---|---|---|---|
| W13 | 1 | Saline | Saline |
| W14 | 1 | Saline | Saline |
| W15 | 3 | Saline | 6mg x 3 |
| W16 | 3 | Saline | 6mg x 3 |
| W17 | 3 | Saline | 6mg x 3 |
| W18 | 3 | Saline | 6mg x 3 |
| W19 | 1 | Saline | Saline |
| W20 | 1 | Saline | Saline |
| W21 | 1 | Saline | Saline |
| W22 | 1 | Saline | Saline |
| W23 | 3 | Saline | 6mg x 3 |
| W24 | 3 | Saline | 6mg x 3 |
| W25 | 3 | Saline | 6mg x 3 |
